# Novel neonatal hypoxic-ischemic model demonstrates neuroinflammation-associated memory deficits without neuronal loss

**DOI:** 10.64898/2026.03.19.712953

**Authors:** Kaylin M. Langer, Erika Tiemeier, Elisabeth Harmon, April Fineberg, Jamie Henry, Isobella Veitch, Tessa Klopper, Tanner McVey, Robert M. Dietz, Andra L. Dingman, Nidia Quillinan

**Author notes:** Co-senior authors (Robert M. Dietz, Andra L. Dingman, Nidia Quillinan). **Corresponding author:** Nidia Quillinan, 12801 E. 17^th^ Avenue, Aurora CO 80045.

## Abstract

**Background:** Neonatal global hypoxic-ischemic cerebral injury is a leading cause of infant mortality and lifelong disability. Current rodent models do not replicate neonatal global cerebral ischemia (nGCI) and reperfusion injury. Here, we developed and characterized a rodent model of cardiac arrest and cardiopulmonary reperfusion (CA/CPR) to induce nGCI, producing acute systemic ischemia, mild neuronal injury, white matter alterations, and motor and memory deficits.

**Methods:** Rat pups underwent CA/CPR or sham procedure on postnatal day 9-11. CA/CPR in rat pups was performed under anesthesia while intubated. Asystole was induced with intravenous (IV) KCl and maintained for 10-14 minutes. Resuscitation included oxygen ventilation, chest compressions, and IV epinephrine.

**Results:** Twelve minutes of asystole provided an optimal balance between survival and systemic injury. Behavioral testing on postoperative day (POD) 7 revealed memory impairments. Despite the absence of overt neuronal death in the hippocampus or cerebellum, we observed evidence of glial activation and white matter alterations.

**Conclusion:** This novel rodent model of nGCI addresses limitations in existing models while offering clinically relevant features to support future mechanistic and translational research.

**Impact:**

- This study validates cardiac arrest and cardiopulmonary resuscitation (CA/CPR) as a novel model for neonatal global cerebral ischemia (nGCI), complementing existing rodent models of unilateral and permanent injury by enabling investigation of both global ischemia and reperfusion injury.
- nGCI results in memory impairment in the absence of overt neuronal cell death. Functional deficits are associated with neuroinflammatory responses in the hippocampus, white matter, and cerebellum.
- Neonatal CA/CPR induces global cerebral ischemia which uniquely allows investigation of hindbrain structures, such as cerebellum, which are typically spared in existing rodent models of neonatal hypoxia-ischemia.

## INTRODUCTION

Neonatal hypoxic ischemia is a leading cause of acute mortality in newborns and long-term neurological morbidity in survivors. Neonatal hypoxic-ischemic encephalopathy (HIE) results from a transient loss of oxygen and blood flow to the brain in the perinatal period ^1^. HIE impacts 1 to 8 per 1000 live births in developed countries and is responsible for ∼23% of neonatal mortality globally^2^. Examples of severe neurological morbidity include cerebral palsy, seizure disorders, cognitive delay, and psychiatric illness^3,4^. Despite advances in neonatal care, treatment options remain limited in application and efficacy. Therapeutic hypothermia (TH), the gold standard of treatment since 2005^5^, reduces mortality and the incidence of cerebral palsy and developmental delay^6–9^. However, the timing of TH initiation is critical and must begin within 6 hours of birth, limiting its application and widespread use in neonates with delayed diagnosis^10,11^. Moreover, infants treated with TH still experience long term neurological deficits^6,10,12–14^. These limitations underscore the critical need for more accessible and broadly effective neuroprotective strategies.

Animal models are critical to our mechanistic understanding of HIE and for developing novel therapies. The most widely used rodent model for neonatal HIE is the Vannucci model^15^, which combines permanent, unilateral carotid artery ligation with systemic hypoxia to produce a unilateral forebrain infarct^16^. This model has been foundational in defining mechanisms of HI injury and has led to important clinical trials and influenced clinical practice^17^. A key advantage of the Vannucci model is that the uninjured contralateral hemisphere serves as an internal control. However, the Vannucci model does not recapitulate global cerebral ischemia or vascular reperfusion injury. Carotid artery occlusion produces a predominantly forebrain-restricted injury, and does not affect the hindbrain, limiting the ability to investigate injury to the cerebellum, an area vulnerable to neonatal HI^1,18,19^. Further, because the vascular occlusion is permanent, the model does not provide insights into vascular reperfusion injury which is critical to the pathophysiology of HIE. While the initial insult depletes the tissue of energy stores, causing acidosis and excitotoxicity, subsequent cerebral reperfusion induces the added physical insult of oxidative stress and metabolic overload, enhancing the overall injury^20^. These model limitations may restrict its translational application^21^. While these limitations have been addressed with larger animal models, such as piglet, ferret, and sheep models^22^, there remains a need for rodent models that produce global injury to enable high-throughput mechanistic and long-term behavioral assessment.

Cardiac arrest and cardiopulmonary resuscitation (CA/CPR) rodent models have been used extensively to study brain injury across the lifespan in rodents as young as PND14^23^, but have not been applied at (PND9-11)^23–27^, which corresponds to the human term-equivalent brain^28^. Here we developed and characterized a novel rodent CA/CPR model of global cerebral ischemia that recapitulates both systemic ischemic injury and central nervous system pathology, including inflammation, white matter alterations, and memory impairment, while sparing gross neuronal death. This work serves as a complement to the work done using the Vannucci model, which is associated with clear histologic cell death. Importantly, up to 63% of children who survive HIE without motor deficits or overt MRI findings are at increased risk of cognitive impairment and behavioral difficulties later in life^29–31^. Accordingly, this model provides a clinically relevant platform to define mechanisms of reperfusion injury and testing therapeutic strategies applicable to a broader range of HIE severity.

## METHODS

All experimental protocols were approved by the Institutional Animal Care and Use Committee (IACUC) at the University of Colorado Anschutz Medical Campus in accordance with the National Institutes of Health guidelines for the care and use of animals in research. Experiments were performed in accordance with ARRIVE 2.0 guidelines^32^. Male and female Sprague Dawley postnatal day 9-11 rat pups from the same litter were randomly assigned to sham or CA/CPR groups. The animals were housed with their dams in a standard 14:10 light: dark cycle and had free access to food and water. Rat pups were randomly assigned to experimental groups. Experimenters performing data collection and analysis were blinded to treatment groups. To minimize potential litter biases, experimental animals were collected from multiple litters. For each behavioral or histologic measure there was no more than n=3/group collected from a single litter and a total of 17 litters were used for behavior and immunohistochemistry experiments.

### Neonatal cardiac arrest with cardiopulmonary resuscitation surgical rat model

Male and female rat pups were subjected to either cardiac arrest and cardiopulmonary resuscitation (CA/CPR) or sham procedures between postnatal day (PND) 9-11. The dam was removed from the home cage and pups were transferred to a surgical procedure room. Rat pups were anesthetized with 3-4% isoflurane, endotracheally intubated with a 24G catheter, and connected to a mini-ventilator set to 160 breaths per minute. Body and pericranial temperature were measured using rectal and ear probes. Normothermia was maintained throughout arrest and resuscitation using a heat lamp to sustain body temperature at 35.5°C±0.3 and a water-filled head coil to sustain pericranial temperature at 37.5°C±0.3°C. A PE-10 catheter was placed in the right internal jugular vein and flushed with heparinized saline for drug administration. Cardiac function was monitored via electrocardiography by placing needle electrodes subcutaneously on the chest. Isoflurane anesthesia was stopped just prior to induction of cardiac arrest. Asystolic cardiac arrest was induced by 50-100µL KCl and injection via a jugular catheter and confirmed on EKG monitor. CPR began 12- or 14-min after induction of cardiac arrest, by reconnecting to the mini-ventilator with delivery of 100% oxygen at a respiratory rate up to 200 breaths/min along with chest compressions and slow injection of 1ml of epinephrine with calcium chloride (0.16 μg epinephrine/ml and 16mg/mL calcium chloride in 0.9% saline). To ensure consistency in total ischemia time, if the return of spontaneous circulation could not be achieved within 3 min of CPR, resuscitation was terminated, and the rat pup was excluded from the study. Following ROSC, animals were weaned from the ventilator and extubated once they achieved a spontaneous breathing rate of 60 breaths per minute. After disconnection, pups received topical lidocaine for analgesia. Shams underwent anesthesia, jugular access, intubation, temperature control, EKG monitoring, and analgesia. Shams did not receive KCl, epinephrine, or chest compressions. After surgical procedures, pups were placed in a clean static cage set on a warming pad. Once surgeries for a given litter were complete, the pups were returned to the home cage and rubbed gently with bedding materials. The dam was then returned to the cage and monitored for 30 minutes to observe for signs of maternal rejection. Postoperatively, pups were visually monitored two times a day for seven consecutive days. Visual monitoring evaluated whether pups sustained necrotic injury and whether they were nested with dam. Topical 2% nitroglycerine ointment was applied to necrosis-affected areas daily. Pups remained in a static cage on a warming pad for three days following the procedure. On day 4, pups and dam were transferred back into their home cage and removed from the heating pad.

Descriptions of methods for all analyses performed after nGCI are reported in the *Supplementary Methods*.

All statistical analyses were performed using GraphPad Prism (Version 10.2.3) unless otherwise stated and are presented as arithmetic mean±standard deviation (SD). For all studies, significance was assigned a p-value of less than 0.05. For two group comparisons, an unpaired two-tailed t-test was used. For comparison of groups with two variables, a 2-way ANOVA with Sidak’s multiple comparisons test was used unless otherwise stated.

## RESULTS

We adapted our previously established CA/CPR model of GCI used in adult and juvenile mice^25,26^ to postnatal day 10 (PND9-11) Sprague-Dawley rat pups, a developmental stage equivalent to full-term human neonates (Figure 1A)^28^. In prior studies, we used 6-8 minutes of asystole in adult mice to induce neuronal injury and approximately 12 minutes in juvenile rats (PND17)^33–35^. In the current model, asystole was induced with intravenous KCl and resuscitation was initiated 12 minutes later. Using this protocol, we successfully resuscitated and achieved survival in 58% of neonates subjected to CA/CPR. Detailed surgical parameters are reported in Table 1. To assess the extent of systemic ischemic injury following resuscitation, we analyzed blood chemistry markers. Serum troponin levels were elevated following GCI, compared to sham controls (8.09±3.91 ng/ml vs. 1.173±0.76 ng/ml, *p*=0.040)(Figure 1B), indicating cardiac ischemia and confirming the presence of multi-organ injury typical of neonates with HIE^36–38^. Additionally, GCI animals exhibited significant acidosis (pH 6.80±0.2 in GCI vs 7.33±0.09 in sham, *p*=0.0029) and elevated serum lactate levels (10.30±2.45 mMol/L in GCI vs 2.47±0.77 mMol/L in sham, *p*=0.0062) (Figure 1C, D). While these markers are not specific for HIE, they are consistent with an ischemic event and are integral to screening for neonatal HIE^39^.

**Figure 1.**
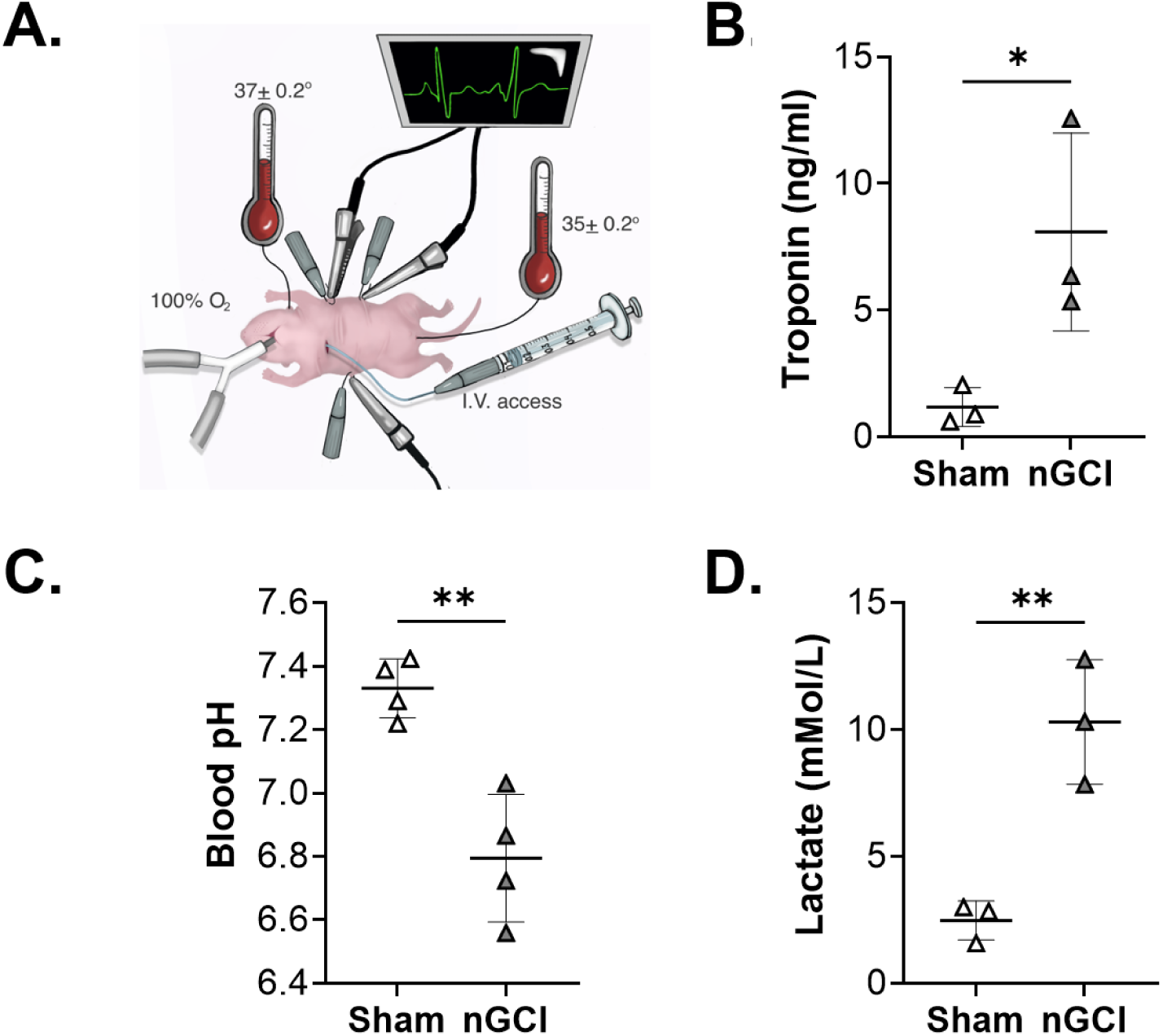
Neonatal rat CA/CPR induces systemic hypoxic-ischemic injury. A.) Graphic representation of neonatal rat CA/CPR model. B) Blood troponin levels (ng/ml) following 12 min CA/CPR animals relative to sham. Whole blood was collected from the right ventricle 2 hours after return of spontaneous circulation. C) Blood pH and D) lactate (mMol/L) following 12 min asystole relative to sham. Whole blood was collected from the right ventricle 20 minutes after return of spontaneous circulation. All animals were sacrificed upon exsanguination. All comparisons analyzed using unpaired t tests (\**p*< 0.05; \*\**p*<0.01). Data represent mean±standard deviation.

**Table 1.**

|  | Sham | nGCI |
| --- | --- | --- |
| N (litters); n (pups) | N=9; n=18 | N=11; n=19 |
| Sex distribution | 9 males, 9 females | 9 males, 10 females |
| Body weight (g) | 18.8 ± 3.3 | 19.3 ± 1.7 |
| Total anesthesia duration (min) | 9.8 ± 2.4 | 10.8 ± 2.4 |
| Total ischemia duration (min) | NA | 13.1 ± 0.5 |
| Total epinephrine for resuscitation (ug) | NA | 0.07 ± 0.02 |
| Return of spontaneous respiration (min) | NA | 16.4 ± <0.001 |
| Total time of intubation (min) | NA | 24.5 ± 0.002 |

Data represent animals included in IHC studies. Data are presented as mean±SD. Total ischemia duration = asystole + resuscitation (concludes upon return of spontaneous circulation). nGCI, neonatal Global Cerebral Ischemia. No differences found among groups.

To evaluate functional consequences of GCI in the neonatal brain, we assessed hippocampal dependent learning and memory. Building on prior work demonstrating hippocampal deficits after GCI in adult and juvenile rodents^40–42^, we tested neonatal rats using contextual fear conditioning 7 days after injury (Figure 2A). nGCI animals exhibited significant reduction in freezing behavior in response to the conditioned context compared to sham controls (16.82%±11.85 nGCI vs. 39.68%±25.57 sham, *p*=0.024) indicating impaired contextual memory. To rule out confounding factors such as locomotor impairment or hyperactivity, we assessed total difference traveled and found no differences between groups (14.82%±9.72 in nGCI,vs.13.48%±4.09 in shams, *p*=0.69)(Figure 2B). Additionally, there were no differences in percentage of time spent in the inner zone of the open field arena, suggesting that anxiety-like behavior did not differ between groups (4.66%±2.2 in nGCI vs. 10.21%±8.05 in sham, p=0.26)(Figure 2B).

**Fig 2.**
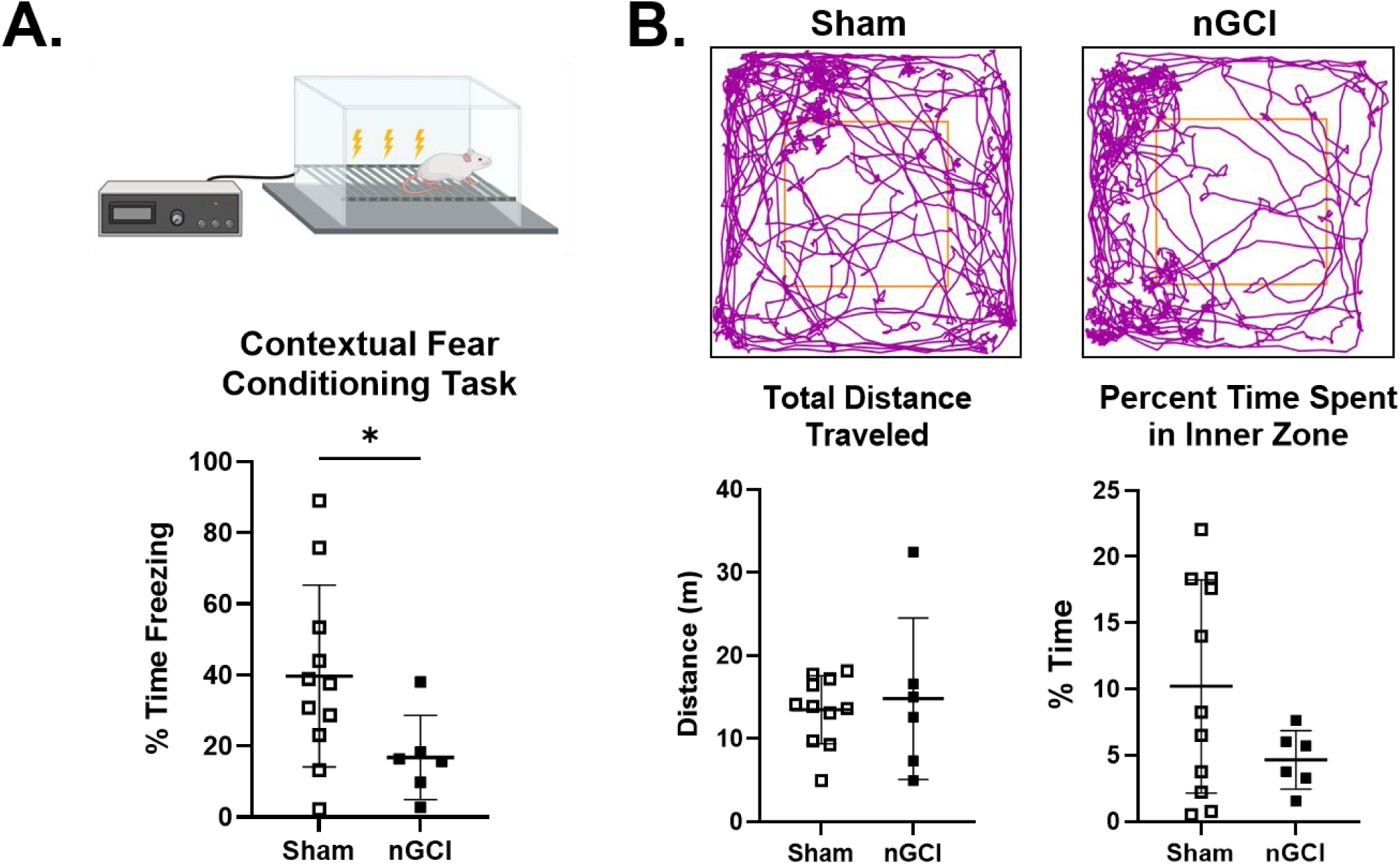
Learning and memory deficits are present after nGCI. A.) Graphic representation of Contextual Fear Conditioning Task (top) and percent time spent freezing 7 days post-nGCI (bottom). B.) Representative track plots in open field test for sham and nGCI (top). Quantified total distanced traveled as well as percentage of time spent in the inner zone of the open field was calculated using ANY-Maze software (bottom). All data compared with Student’s t-test; \**p*< 0.05. Data represent mean±standard deviation. Sham: N (litters)=6, n (pups)=11; males=6, females=5. nGCI: N=3, n=6; males=3, females=3.

In light of the memory impairments observed in our model, we next evaluated whether nGCI induced acute loss of hippocampal neurons. Fluorojade C (FJC) staining was performed in hippocampal brain sections to fluorescently label dead and dying neurons (Figure 3A, B). Surprisingly, no FJC-positive cells were detected in the hippocampus at 3- or 7- days after injury (3 days not shown). The absence of FJC staining is unlikely to reflect technical failures, as FJC labeling was observed in neonates subjected to a longer duration of asystole (14 minutes), suggesting that neuronal death was not induced with the 12-minute protocol. While this shows lack of active cell death at 7 days, to rule out earlier cell death, we conducted a neuronal density analysis in CA1 and CA3 hippocampal subregions. No differences were observed between nGCI and sham animals, (CA1: 0.44±0.073 cells/mm^3^ in nGCI vs. 0.44±0.026 cells/mm^3^ in sham, *p*=0.99; CA3: 0.28±0.045 cells/mm^3^ in nGCI vs. 0.28±0.029 cells/mm^3^ in sham, *p*=0.71)(Figure 3C), consistent with our FJC staining. These data reveal a notable dissociation between functional memory impairments and overt hippocampal neuron death following 12 minutes of asystole.

**Fig 3.**
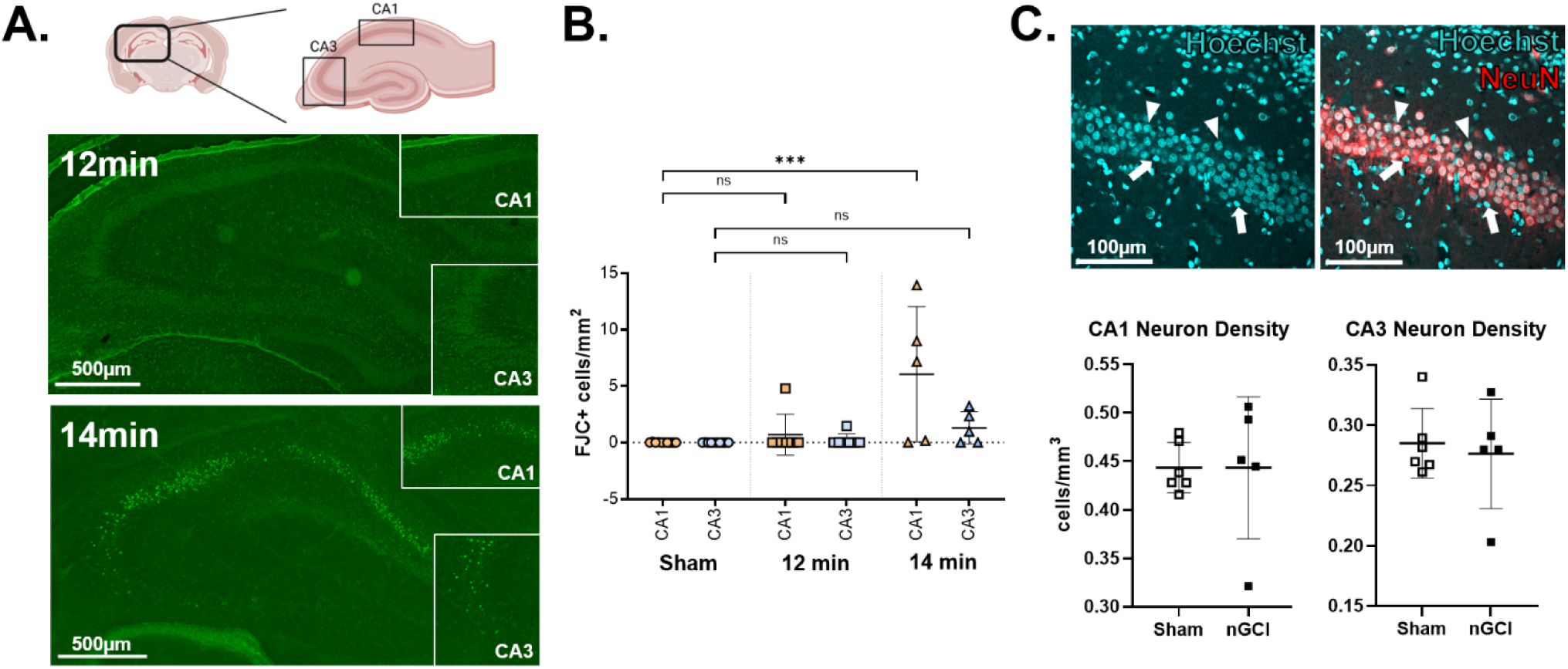
Resiliency of the neonatal brain to GCI. A.) Graphical representation of analyzed hippocampal subregions, CA1 and CA3 (top) and representative Fluorojade C staining of hippocampus at 7 days post-nGCI following 12 or 14 minutes of cardiac arrest (bottom). B.) Quantification of Fluorojade C positive cells in CA1 and CA3 Hp subregions across groups. One-way ANOVA followed by a post hoc Tukey’s test for multiple comparisons (\*\*\**p*<0.001). C.) Representative Hoechst and NeuN staining (top) and quantification of live neuronal densities (bottom) 7 days post-nGCI. Student’s unpaired t-test. Data represent mean±standard deviation.

Given the absence of overt neuronal cell death despite clear deficits in hippocampal-dependent memory following 12 minutes of asystole, we next investigated whether neuroinflammation might contribute to hippocampal dysfunction following nGCI. Microglia and astrocytes play central roles in modulating injury responses and regulating synaptic plasticity that underlies hippocampal dependent memory after ischemia^43–49^. Therefore, we assessed glial activation in the CA1 and CA3 subregions. At 7 days post-resuscitation, immunohistochemical analysis revealed significantly increased Iba1 immunoreactivity in GCI animals compared to sham controls, staining (CA1: 30.48%±2.73 in nGCI vs. 21.86%±4.29 in sham, *p*=0.004; CA3: 19.98±4.06 in nGCI vs. 15.53±2.28 in sham, *p*=0.047) consistent with microglial activation (Figure 4). Similarly, GFAP staining was markedly elevated in both subregions following nGCI, reflecting reactive astrogliosis (CA1: 19.84%±2.04 in nGCI vs. 10.34%±5.66 in sham, *p*=0.006; CA3: 21.86±3.04 in nGCI vs. 11.89±6.67 in sham, *p*=0.013). Together, these findings reveal a robust and spatially broad glial response in the hippocampus, despite preserved neuronal structure. This neuroinflammatory signature may contribute to the observed hippocampal memory impairments, even in the absence of overt neurodegeneration.

**Fig 4.**
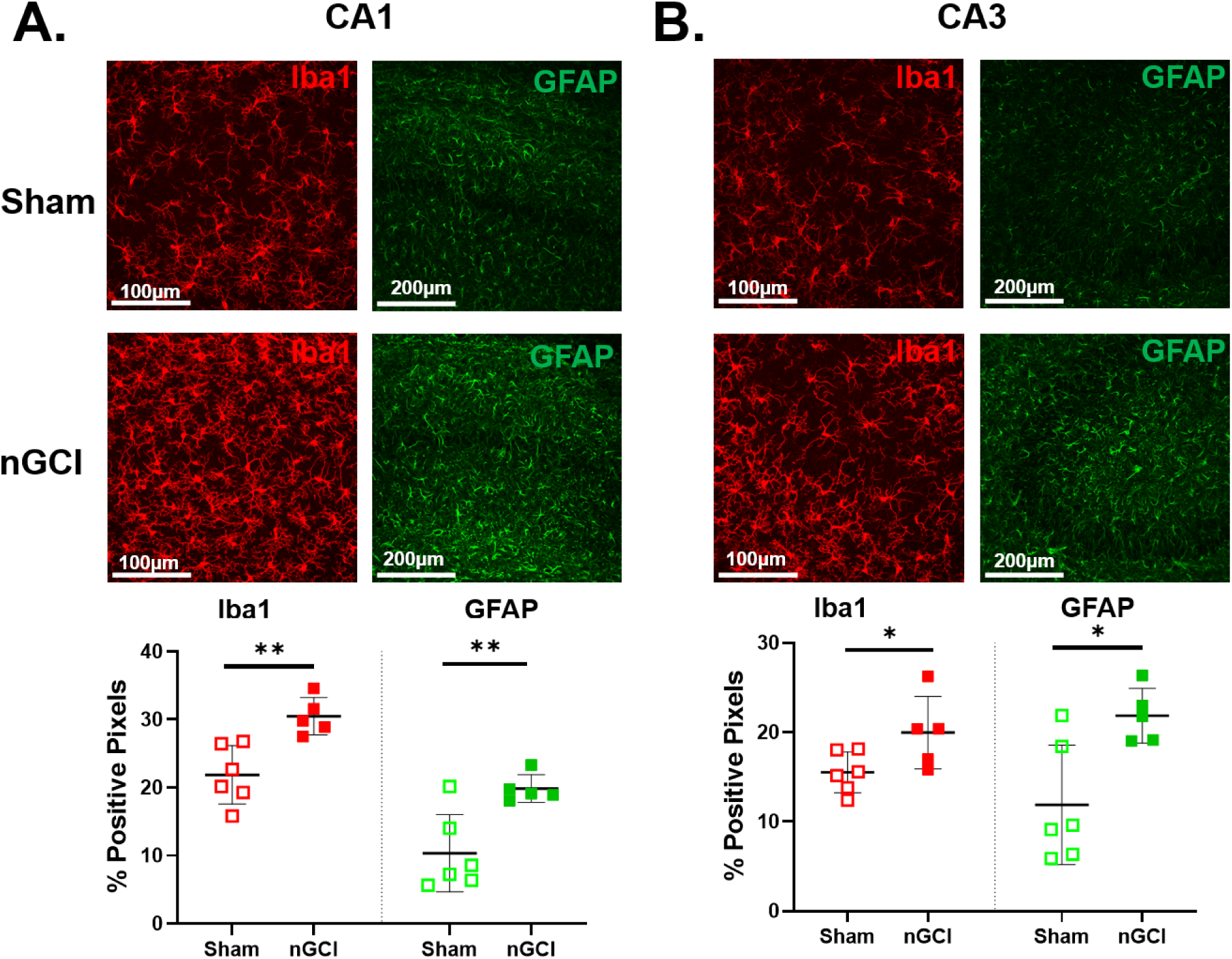
Hippocampal neuroinflammation 7 days after nGCI. A.) Iba1 and GFAP immunohistochemistry representatives and quantification of the CA1 region. b. Iba1 and GFAP immunohistochemistry representatives and quantification of the CA3 region. All comparisons analyzed using Student’s unpaired t tests. \**p*< 0.05; \*\**p*<0.01. Data represent mean±standard deviation.

White matter alterations are commonly observed following HIE^4,50,51^ and are associated with long-term impairments. To determine whether nGCI impacts white matter microstructure we focused on the corpus callosum (CC), the largest white matter tract in the forebrain. Myelin structure and lipid content were assessed using spectral microscopy, with polarity index serving as an inverse measure of cholesterol content and myelin integrity^52,53^. Medial and lateral subregions of the CC were imaged separately at 3- and 7-days post-resuscitation. At 3 days, spectral emission signatures corresponding to myelin were undetectable in the medial CC, precluding reliable polarity index analysis in this region. In contrast, the lateral CC retained detectable myelin signal, and polarity index measurements revealed a significant increase in polarity index at 7 days post-injury in nGCI compared to sham (0.52±0.06 in sham vs 0.62±0.03 in 7-day nGCI, *p*=0.021)(Figure 5A), suggesting compromised myelin microstructure due to injury. Additionally, the polarity index increased between 3- and 7-days post-nGCI in the lateral CC(0.50±0.07 in 3-day nGCI, *p*=0.008) indicating a delayed and progressive alteration of the white matter microstructure.

**Fig 5.**
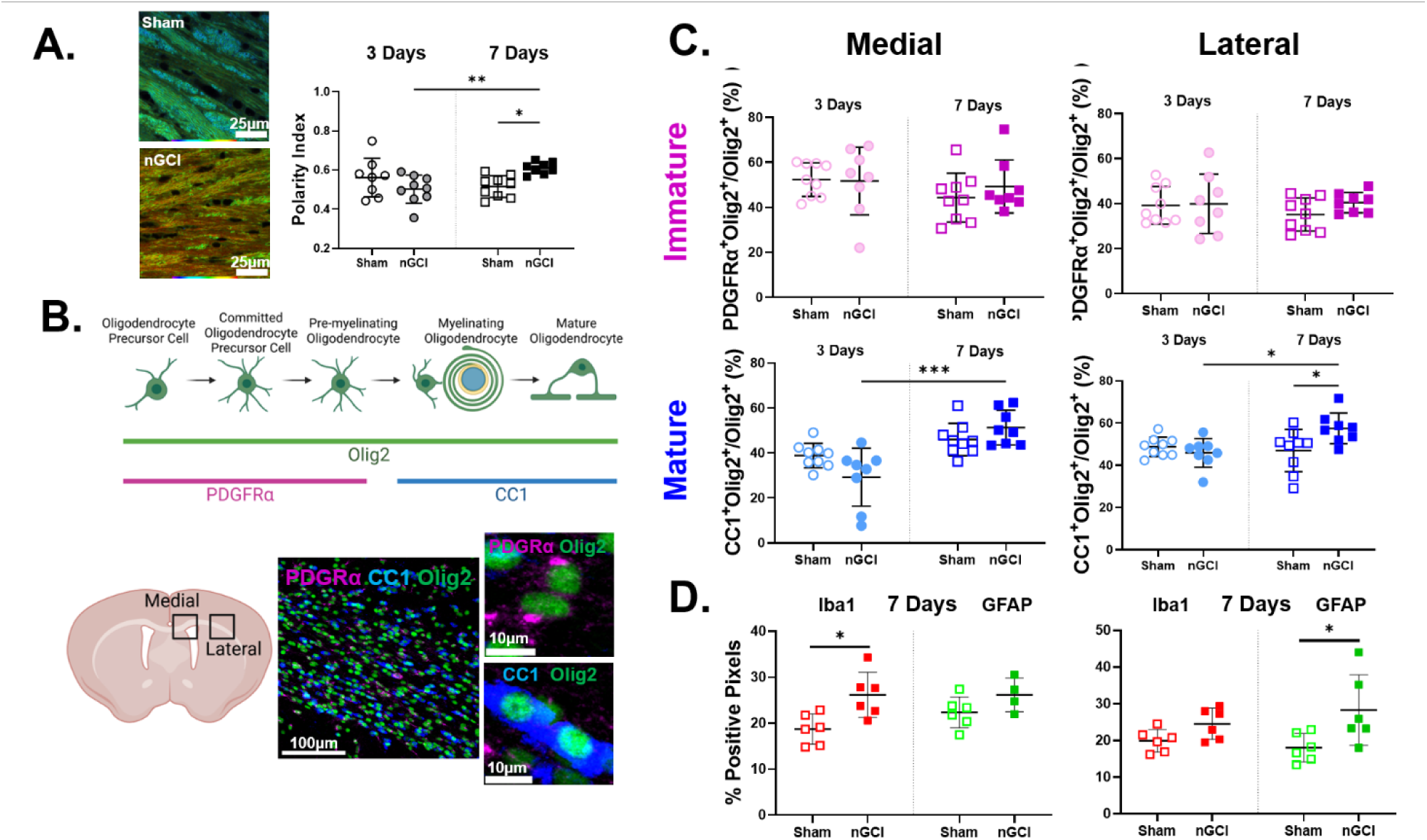
White matter microstructure changes in corpus callosum (CC). A.) Representative Nile Red spectral microscopy images with pseudocoloring and quantification of polarity index in the CC at 3 and 7 days post-nGCI (left). Polarity index for each pixel was calculated as the ratio of the maximum intensity wavelength to the full spectral range (right). B. Schematic of oligodendrocyte (OL) maturation stages with corresponding immunohistochemistry (top). Graphic of analyzed CC regions (medial and lateral) and representative immunohistochemistry images (bottom). C.) Quantification of the proportion of immature OLs (PDGFRα^+^Olig2^+^/Olig2^+^) in medial and lateral CC (top), mature OLs (CC1^+^Olig2^+^/Olig2^+^) (bottom) D.) Iba1 and GFAP immunohistochemistry quantification 7 days post-nGCI in medial (left) and lateral (right) CC. All comparisons analyzed using two-way ANOVA followed by Sidak’s multiple comparison test (\**p*< 0.05; \*\**p*<0.01; \*\*\**p*<0.001). Data represent mean±standard deviation.

To investigate potential cellular mechanisms underlying these white matter changes, we analyzed oligodendrocyte maturation using immunohistochemistry. Oligodendrocyte progenitor cells (OPCs) were defined as Olig2^+^ cells co-expressing PDGFRα (PDGRFα^+^Olig2^+^/Olig2^+^), while mature and myelinating oligodendrocytes as those co-expressing Olig2 and CC1(CC1^+^Olig2^+^/Olig2^+^)^54^. Given developmental differences in myelination across CC subregions^55^, medial and lateral CC were analyzed separately at 3- and 7-days after nGCI (Figure 5B). Cell counts were performed to identify a potential shift in oligodendrocyte differentiation(Figure 5C). At 3 days, there were no significant differences in the proportions of immature or mature oligodendrocytes between nGCI and sham animals. However, by 7 days post-injury, nGCI animals showed an increase in the proportion of mature oligodendrocytes compared to sham in the lateral CC (46.98%±9.98 in sham vs. 57.52%±7.26 in nGCI, p=0.027). Notably, both the medial and lateral CC regions demonstrated an increase in mature oligodendrocytes between 3- and 7-days post-injury (medial: 29.22%±12.87 in 3-day nGCI vs. 51.28%±7.74 in 7-day nGCI, p=0.004; lateral: 45.89%±6.82 in 3-day nGCI vs. 57.52%±7.26 in 7-day nGCI, p=0.02), suggesting an acceleration of the maturation of oligodendrocytes into myelinating cells following ischemia. Finally, to explore the contribution of gliosis to changes in OL maturation and white matter microstructure at 7 days post-nGCI, we again assessed glial activation in the medial and lateral subregions. Immunohistochemical analysis revealed significantly increased Iba1 immunoreactivity in GCI animals compared to sham controls in the medial CC only (Medial: 26.13%±4.91 in nGCI vs. 18.73%±3.30 in sham, *p*=0.012; Lateral: 24.54±4.21 in nGCI vs. 19.92±3.07 in sham, *p*=0.055) consistent with microglial activation (Figure 5). Analysis showed increased GFAP immunoreactivity in only the lateral CC in GCI animals relative to shams (Medial: 26.15%±3.66 in nGCI vs. 22.36%±3.37 in sham, *p*=0.13; Lateral: 28.27±9.57 in nGCI vs. 18.01±3.86 in sham, *p*=0.035)(Figure 5).

Given our model’s ability to assess both forebrain and hindbrain structures, we next investigated whether the cerebellum, an area vulnerable to hypoxic-ischemic injury, exhibited histopathologic (cell death and inflammation) changes following nGCI. We first focused on Purkinje cells in the cerebellar cortex. These large neurons are critical for motor coordination and highly sensitive to ischemic injury. At 7 days post-injury, there were no significant differences in Purkinje cell density between nGCI and sham animals (0.262±0.08 cells/mm^2^ in nGCI vs. 0.257±0.04 cells/mm^2^ in sham, *p*=0.88)(Figure 6A), mirroring the absence of cell loss in the hippocampus. We next examined glial activation within the molecular layer, which contains the expansive dendritic arbors of Purkinje neurons. Immunostaining revealed significant increases in both Iba1 and GFAP signal in nGCI animals compared to sham (Iba1: 8.79%±1.98 in nGCI vs. 6.41%±1.36 in sham, *p*=0.019; GFAP: 35.04%±4.86 in nGCI vs. 28.78±4.92 in sham, *p*=0.024)(Figure 6B,C), indicating microglial and astrocyte activation in this region. To assess cerebellar white matter integrity, we again used spectral microscopy to quantify polarity index. At 7 days post-injury, nGCI animals demonstrated a significant increase in polarity index compared to sham controls (0.67±0.09 in nGCI vs. 0.57±0.07 in sham, *p*=0.014)(Figure 6D), suggesting disrupted white matter microstructure and altered lipid organization similar to that observed in the lateral corpus callosum. However, in contrast to the corpus callosum, we did not observe corresponding increases in oligodendrocyte maturation (Immature: 31.82%±5.8 in nGCI vs. 28.76%±3.6 in sham, *p*=0.22; Mature: 54.27%±12.3 in nGCI vs. 54.75±10.75 in sham, *p*=0.93)(Figure 6E), nor did we detect differences in Iba1 or GFAP staining in the cerebellar white matter (Iba1: 22.94%±5.51 in nGCI vs. 19.20%±7.52 in sham, *p*=0.34; GFAP: 38.43%±6.28 in nGCI vs. 35.34±5.25 in sham, *p*=0.32)(Figure 6F). These findings suggest that white matter injury in the cerebellum may occur independently of local glial activation and through distinct mechanisms compared to the forebrain.

**Fig 6.**
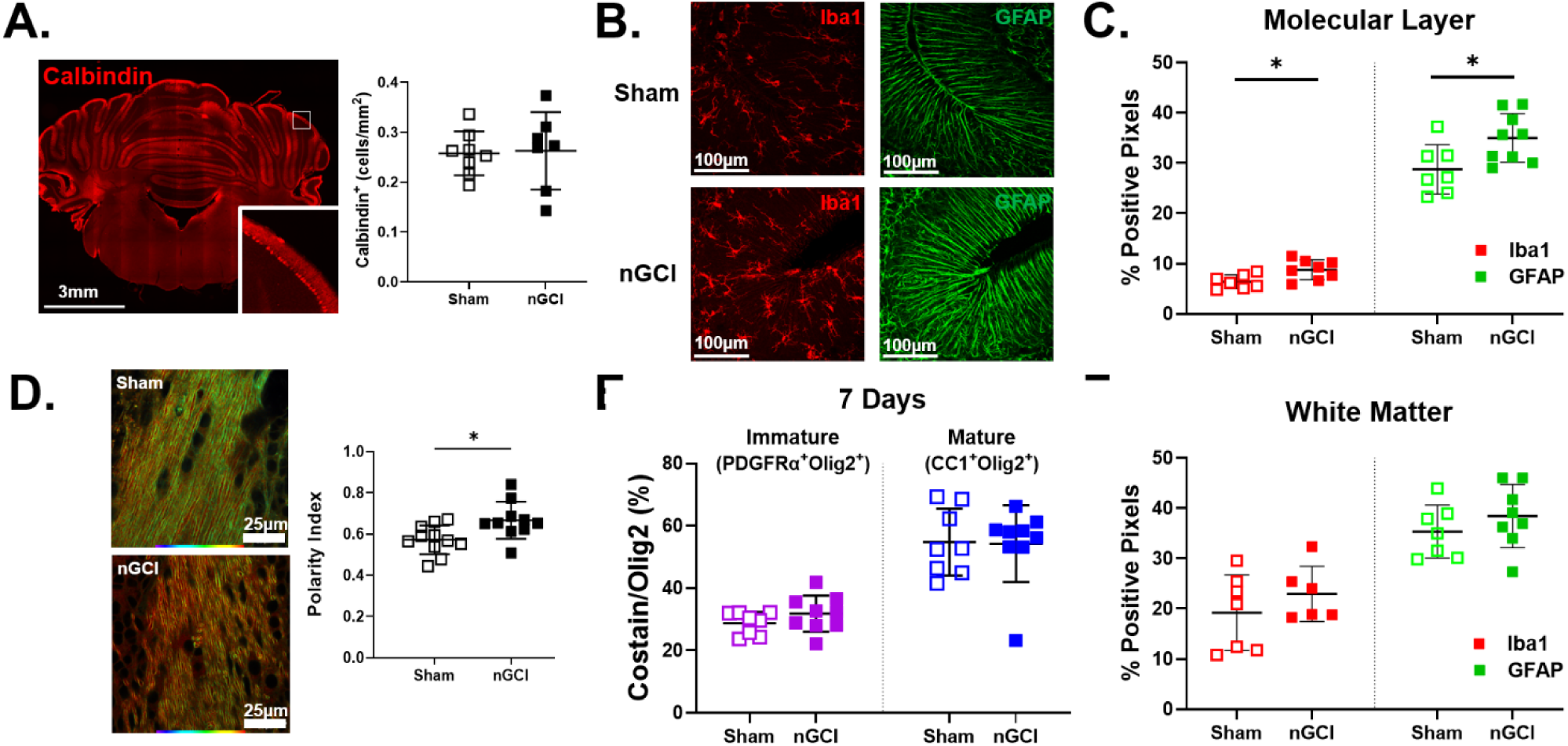
Gliosis and white matter abnormalities in the cerebellum 7 days following nGCI. A.) Representative image of the cerebellum calbindin staining to identify Purkinje cells and quantification of Purkinje cell density. B.) Representative images of Iba1 and GFAP immunohistochemistry and C.) quantification in the molecular layer of the cerebellum. D.) Representative images of Nile Red spectral microscopy in the cerebellar white matter with pseudocoloring and quantification of polarity index 7 days post-nGCI. E.) Quantification of the proportion of immature OLs (PDGFRα^+^Olig2^+^/Olig2^+^) and mature OLs (CC1^+^Olig2^+^/Olig2^+^) in the cerebellar white matter. F.) Quantification of Iba1 and GFAP immunohistochemistry in the cerebellar white matter. All comparisons analyzed using unpaired t-tests (\**p*< 0.05). Data represent mean±standard deviation.

## DISCUSSION

In this study, we developed and characterized a novel rodent model of neonatal CA/CPR that, to our knowledge, is the first to model global cerebral ischemia with reperfusion injury in term-equivalent neonates. This model produces learning and memory deficits at 7 days post-injury despite resilience to cell death. In the forebrain, nGCI induced microglia and astrocyte activation in the CA1 and CA3 subfields of the hippocampus, along with white matter disruption in the corpus callosum at 7 days post-injury. White matter alterations were further characterized by accelerated oligodendrocyte differentiation, as evidenced by increased proportions of mature oligodendrocytes 7 days after injury. Importantly, this model also allows for the study of hindbrain injury, which is underrepresented in traditional models of hypoxia-ischemia. In the cerebellum, we observed a similar pattern of resilience to cell death and activation of glial cells within the molecular layer, but with distinct dissociation in the white matter. Despite altered white matter microstructure, there were no changes in oligodendrocyte differentiation or glial activation. These findings underscore the regional specificity of neonatal brain responses to ischemic injury and highlight the importance of evaluating hindbrain and forebrain separately. Extrapolating mechanisms from across regions may obscure critical differences in vulnerability, temporal evolution, and repair processes in the developing brain. This study provides essential histological data that will serve as a critical foundation for future mechanistic and hypothesis-driven studies of global neonatal ischemia.

The ability of MRI to predict neurodevelopmental outcomes in neonates with global cerebral hypoxic-ischemic injury is limited, particularly among infants without detectable injury. Notably, hippocampal injury in neonates with HIE often escapes early MRI detection^56^ yet is strongly associated with long-term cognitive and emotional regulation deficits, even in those treated with therapeutic hypothermia^13,57^. Our model addresses this gap by demonstrating measurable cognitive dysfunction following neonatal hypoxic ischemic injury that is independent of overt neuronal cell death or necrosis. The Vannucci model has a well-established injury pattern of multiple necrotic foci within both gray and white matter^17^ relevant to mechanisms underlying neurological impairments and cerebral palsy^21^. In contrast, our model captures a clinically relevant pattern of hippocampal dysfunction, microglial and astrocytic activation, and white matter alterations, without widespread neurodegeneration, reflecting the presentation of many children who survive without MRI-detectable injury but experience cognitive or psychiatric difficulties later in life^13,29–31,58,59^. These differences likely reflect key distinctions from the Vannucci model, including global vs. unilateral injury and reperfusion vs. permanent injury. Accordingly, this model can be used alongside the Vannucci model to delineate how injury pattern and vascular dynamics (reperfusion vs permanent occlusion) differentially regulate inflammatory and apoptotic cascades. One advantage of the neonatal GCI model described here is the ability to investigate whole-body HI in neonates. While large animal studies remain essential to these investigations, this rodent model provides a high throughput complement for mechanistic and behavioral investigation. Interpretation should be considered in the context of model-specific limitations, including the use of hyperkalemic arrest rather than hypoxic arrest, which more closely models the predominant clinical etiology of HIE, perinatal asphyxia^2^. Additionally, our model does not capture factors unique to the intrapartum environment, reinforcing the role of neonatal rat CA/CPR as a complement to, not replacement of, existing HI models. Rather, our model provides a platform for mechanistic and behavioral studies of neonatal GCI and for identifying strategies to improve long term outcomes.

Nonetheless, the dissociation between cognitive impairment and neuronal cell death in our model is somewhat unexpected given the well-established vulnerability of the hippocampus and cerebellum to GCI in adult models^60–62^. Anesthesia exposure and hypothermic body temperatures may be neuroprotective in cerebral ischemia ^63–65^, raising the possibility that anesthesia exposure prior to arrest modulates overall injury. However, anesthesia in this study is consistent between sham and injured animals and with studies of adult animals where neuronal cell death is observed ^40,66,67^. Similarly, normothermia was maintained, minimizing the possibility that these conditions are the sole driver of neuronal resilience. Moreover, increasing the asystole time to 14 minutes resulted in neuronal cell death, supporting the model’s capacity to induce graded injury and enabling titration across the full spectrum of severity. Previous CA/CPR models in adult rodents have shown hippocampal cell death driven by an initial cascade of energy failure, excitotoxicity, calcium dysregulation, and oxidative stress^20,45,68^, followed by a latent phase (hours to days) characterized by activation of the apoptotic cascade and inflammatory pathways^69,70^. In contrast, our findings align with prior work demonstrating increased resilience to GCI in younger brains.^26^ Asphyxial cardiac arrest models applied in rats ranging from P14-17 with similar ischemia times used here have also demonstrated hippocampal degeneration^23,33,34^ suggesting that the term-equivalent neonatal brain may have a higher threshold for ischemic injury than older age groups, although differences of hypoxia preceding GCI in asphyxial models may contribute. The mechanisms underlying the relative resilience in neonates remain undefined and warrant future investigation, including studies that titrate asystole time to produce varying degrees of neuronal injury.

Importantly, histological findings in the cerebellum mirrored those in the hippocampus, with no Purkinje cell loss but evidence of glial activation and white matter disruption, suggesting a global neuronal resilience to GCI at this developmental stage. A limitation of this study is the relatively narrow time course evaluated. Sub-acute time points for FJC analysis of cell death (3-7 days post-resuscitation) were chosen based on the known temporal evolution of injury in clinical MRI and prior histological studies^26,34,60,67,71–77^, as well as the presence of a clear behavioral deficit at 7 days. Future studies should extend histological and behavioral outcomes to later time points including puberty and adulthood to define the long-term consequences relevant to cognitive and emotional deficits. Chronic behavioral assessments should also be expanded to capture neuromotor, cognitive, and affective domains impacted in survivors of neonatal HIE.

Acute increases in circulating inflammatory markers are observed in neonatal HIE and correlate with neurological outcomes, even in the absence of detectable MRI changes^69,78,79^. In our model, memory deficits were associated with increased glial reactivity, consistent with neuroinflammation, but not with neuronal loss. Increased astrocyte and microglia reactivity were evident in both the hippocampus and cerebellar grey matter, but not cerebellar white matter, suggesting region specific inflammatory responses to GCI. While therapeutic hypothermia is believed to confer neuroprotection in part by attenuating inflammation, the evolution of HIE injury is likely more complex. TH suppresses microglia activation and release of IL-1β, TNF-α, IL-6 which exacerbate neuronal and oligodendrocyte injury^80–83^. Yet, recent data demonstrate persistent microglia inflammatory markers despite reduced neuronal injury following TH^84^, aligning with clinical studies that show almost one third of infants treated with TH develop moderate to severe impairments^6,7^. While microglia activation is thought to contribute to apoptosis and neurodegeneration^34,47^, other studies suggest that microglia also exert protective effects after HIE^85,86^ and their depletion is detrimental^87^. This duality likely reflects the important roles of microglia in normal brain development^88–91^ and phagocytosis of cellular debris^88,92–94^. These studies highlight the heterogeneity in microglial responses and the need to define context-specific activation states following neonatal HI injury. Similarly, we observed increased GFAP expression after nGCI, consistent with reactive astrogliosis, as observed in other HIE studies^84,95,96^. Elevated plasma GFAP levels have been observed in patients after HIE and are correlated with worse neurological outcomes^79,97^, suggesting a potential detrimental role. However, similar to microglia, astrocytes play important roles in development, and a more nuanced understanding of astrocytic responses to GCI is needed. Indeed, studies using cultured astrocytes show that hypothermia enhances the expression of protective factors following simulated ischemia, suggesting potentially beneficial astrocytic functions in the post-ischemic brain^98^. Given the central role of CNS inflammation, future studies should examine the temporal evolution of emerging clinical biomarkers, including inter-alpha inhibitor proteins (IAIPs), neuron-specific enolase (NSE) and S100-calcium-binding protein-B (S100B)^99,100^ to refine diagnostic and mechanistic insight. While the current study supports an association between inflammation and memory deficits, future studies that manipulate inflammatory pathways are necessary to establish causality.

White matter injury is common in models of preterm and term neonatal HI^4,51,101^, and we demonstrate white matter microstructure following nGCI, reflected by changes related to altered cholesterol content both at 7 days and progressing between 3- and 7-days post-injury. We also see increased astrogliosis in forebrain white matter 7 days after nGCI. Oligodendrocytes are myelin producing glial cells within the brain and have been well studied using the Vannucci model, where injury often leads to impaired differentiation and vulnerability of oligodendrocyte precursor cells^102,103^. In contrast, our model showed no changes in immature oligodendrocytes, possibly related to global ischemia, rather than the more local ischemia of the Vannucci model, or developmental timing of injury. While we did not see changes in the OPC lineage, we observed increases in mature, myelinating oligodendrocytes, suggesting that while oligodendrocytes are progressing to a mature phenotype, their capacity to produce or traffic cholesterol may be altered, affecting myelin structure. Cholesterol synthesis is a key component of normal developmental myelination. Membrane cholesterol promotes myelin compaction^104^ and promotes synthesis of myelin proteins such as proteolipid protein^105^. Altered axonal signaling for myelin formation may be also be disrupted^106^. The restriction of findings to the lateral corpus callosum may reflect the known lateral to medial progression of myelin development and maturation in early development^55^. Further characterization of myelin integrity and oligodendrocyte lineages will help to understand the impacts of nGCI on white matter development.

Finally, a unique advantage of our model is its ability to simulate neonatal hypoxic-ischemic injury in the hindbrain, a region frequently underestimated by neonatal MRI despite its known vulnerability^19^. Early cerebellar dysfunction and anatomical abnormalities are increasingly recognized as contributors to impaired cerebral growth and developmental disorders, including autism spectrum disorder, attention deficit hyperactivity disorder, and developmental dyslexia^107^. While forebrain networks develop prenatally, the cerebellum undergoes prolonged postnatal development and plays critical roles for motor coordination and cognition^108–110^ with important cerebellocortical interactions developing over the first 5 years of life^111,112^. This extended developmental window may increase vulnerability to HI that occurs at term birth. In our model, the molecular layer, which houses the dense dendritic fields of Purkinje cells, showed glial activation without cell death, paralleling our findings in the hippocampus. In the cerebellar white matter, we also observed disrupted microstructure without corresponding changes in oligodendrocyte lineage or glial reactivity. Together, these findings suggest distinct mechanisms of white matter injury in the developing cerebellum and highlight the need to study the cerebellum as a separate entity from the forebrain, while reinforcing the utility of this model for dissecting region-specific injury. This study is limited in its assessment of neuromotor outcomes, in part due to rudimentary mobility at these timepoints. Gross motor deficits were not observed, as indicated by no differences in total distance traveled in an open field test, but more sensitive neuromotor tasks such as negative geotaxis and righting reflex would provide a more nuanced assessment of motor function after this injury. Later timepoints will also enable socioemotional testing, particularly given the association between HIE and autism spectrum disorder. Future studies are necessary to more comprehensively define the long term neuromotor and cognitive outcomes following neonatal global cerebral ischemia.

## Conclusions

Neonatal rat CA/CPR provides a novel and clinically relevant model of neonatal global cerebral ischemia with reperfusion injury that recapitulates key features of the pathophysiology of HIE. This model addresses critical gaps in HIE research by enabling high-throughput investigation of reperfusion injury, GCI injury in the hindbrain, and the mechanisms underlying cognitive and behavioral deficits associated with HIE.

